# Bioorthogonal non-canonical amino acid tagging reveals translationally active subpopulations of the cystic fibrosis lung microbiota

**DOI:** 10.1101/588640

**Authors:** Talia D. Valentini, Sarah K. Lucas, Kelsey A. Binder, Lydia C. Cameron, Jason A. Motl, Jordan M. Dunitz, Ryan C. Hunter

**Affiliations:** Department of Microbiology & Immunology, University of Minnesota, 689 23rd Avenue SE, Minneapolis, MN 55455; Academic Health Center, University Flow Cytometry Resource, University of Minnesota, 6th St SE, Minneapolis, MN 55455; Division of Pulmonary, Allergy, Critical Care & Sleep Medicine, University of Minnesota, 420 Delaware St. SE, Minneapolis, MN 55455

## Abstract

Culture-independent studies of cystic fibrosis lung microbiota have provided few mechanistic insights into the polymicrobial basis of disease. Deciphering the specific contributions of individual taxa to CF pathogenesis requires a comprehensive understanding of their *in situ* ecophysiology. We applied bioorthogonal non-canonical amino acid tagging (BONCAT), a ‘click’ chemistry-based metabolic labeling approach, to quantify and visualize translational activity among CF microbiota. Using BONCAT-based fluorescent imaging on sputum collected from stable CF subjects, we reveal that only a subset of bacteria are translationally active. We also combined BONCAT with fluorescent activated cell sorting (FACS) and 16S rRNA gene sequencing to assign taxonomy to the active subpopulation and found that the most dominant taxa are indeed translationally active. On average, only ∼12-18% of bacterial cells were BONCAT labeled, suggesting a heterogeneous growth strategy widely employed by most airway microbiota. Differentiating translationally active populations from those that are dormant adds to our evolving understanding of the polymicrobial basis of CF lung disease and may help guide patient-specific therapeutic strategies targeting active bacterial populations that are most likely to be susceptible.

---

The increased viscosity and impaired clearance of mucus secretions in cystic fibrosis (CF) airways creates a favorable environment for chronic microbial colonization, the primary cause of patient morbidity and mortality (1). *Pseudomonas aeruginosa* and *Staphylococcus aureus* have long been recognized as primary CF pathogens and are the targets of common therapeutic regimens (2), though recent culture-independent studies have revealed a more complex polymicrobial community harboring facultative and obligately anaerobic bacteria that are relatively understudied (3-5). While the specific contributions of individual community members to disease progression remain poorly understood and at times controversial (6), cross-sectional studies of both pediatric and adult cohorts have revealed compelling relationships between bacterial community composition and disease state, antibiotic use, patient age and other phenotypes (7-12). These data have challenged the field to reconsider therapeutic strategies in a polymicrobial community context (13,14).

Relatively fewer studies have identified within-patient perturbations in bacterial community structures that coincide with pulmonary exacerbations (PEx), characterized by increased respiratory symptoms and an acute decrease in lung function. In general, PEx symptoms are resolved in response to antibiotic therapy (validating a bacterial etiology), though sputum cultures generally demonstrate that airway pathogens are recovered at similar densities before, during and after disease flares (15-18). Culture-independent studies show similar trends; with few exceptions (9,19,20) longitudinal sequencing analyses of sputum from individual subjects frequently reveal unique, patient-specific bacterial communities whose diversity and composition remain stable during PEx onset and upon resolution of patient symptoms (15,21,22). This lack of association between lung microbiota and disease dynamics may reflect the inability of both culture-based and sequencing approaches to capture changes in bacterial activity, which likely have a critical impact on disease progression and patient response to therapy.

To date, there have been few studies of bacterial growth and metabolism within the CF airways (23-28). RNA-based profiling of stable CF subjects has shown consistencies between rRNA and rDNA signatures suggesting that many bacterial taxa identified by 16S rRNA gene sequencing are transcriptionally active, though these data have also corroborated that bacterial community membership is not necessarily predictive of *in vivo* growth activity (23,24). Indeed, interactions between respiratory pathogens and the host and/or co-colonizing microbiota can influence growth rates, metabolism, virulence factor production and antimicrobial susceptibility without an accompanying change in bacterial abundance (29-34). Moreover, growth rates of respiratory pathogens can vary substantially between subjects and even within a single sputum sample (25,26), the heterogeneity of which is not captured using conventional molecular profiling. There remains a need for novel methods to characterize *in situ* bacterial activity and its role in disease progression.

Bioorthogonal non-canonical amino acid tagging (BONCAT) has been used to characterize the activity of uncultured microbes in soil and marine samples (35-38). BONCAT relies on the cellular uptake of a non-canonical amino acid (*e.g.* L-azidohomoalanine (AHA), a L-methionine analog) carrying a chemically-modifiable azide group. After uptake, AHA exploits the substrate promiscuity of methionyl-tRNA synthetase and is incorporated into newly synthesized proteins. Translationally active cells can then be identified through a bioorthogonal azide-alkyne ‘click’ reaction in which a fluorophore-tagged alkyne is covalently ligated to AHA, resulting in a fluorescently labeled population of cells that can be further studied using a variety of microscopy and analytical methods. BONCAT has been shown to correlate with other established methods of quantifying microbial activity (36) and represents a robust tool for characterization of bacterial communities in their native growth environment.

BONCAT has also recently been used to study bacterial pathogens *in vitro* (39-42), though it has seen limited use in the study of human host-associated bacterial communities (36). Samples derived from the CF airways provide a unique opportunity to do so, as the site of infection is amenable to longitudinal studies and the bacterial growth environment is relatively stable upon removal from the host (43). Exploiting these advantages, we used BONCAT together with fluorescence-activated cell sorting (FACS) and 16S rRNA gene sequencing to characterize the *in situ* translational activity of bacterial communities within sputum. We reveal that active bacteria represent only a subset of microbiota captured using conventional 16S rRNA gene sequencing and discuss these results in the context of airway disease progression and treatment of individual patients.

## RESULTS

### BONCAT differentiates translationally active and inactive *P. aeruginosa* cells *in vitro*

To optimize the BONCAT experimental approach, we first grew *P. aeruginosa*, a canonical CF pathogen, to mid-log phase followed by supplementation with 6mM L-azidohomoalanine (AHA) for 3h. Post-AHA treatment, azide-alkyne ‘click chemistry’ using Cy5 labeled dibenzocyclooctyne (Cy5-DBCO) permitted fluorescent detection of translationally-active cells (Fig. 1a). Quantification of average Cy5 pixel intensity per cell revealed active protein synthesis in ∼98% of the population. By contrast, supplementation of the growth medium with 6mM L-methionine (MET) or pre-treatment of *P. aeruginosa* with tobramycin, chloramphenicol and tetracycline (to arrest *de novo* protein synthesis) prior to AHA resulted in negligible fluorescence (Fig. 1b,c). These data were also confirmed by SDS-PAGE (Supplementary Fig. 2). Finally, when two AHA labeled cultures (one treated with antibiotics, one without) were combined in a 1:1 ratio prior to Cy5-DBCO labeling, two subpopulations with only minor overlap in fluorescence intensity were identified, representing a mix of active and inactive cells (Fig. 1d). Together, these data demonstrate the utility of BONCAT for characterizing *P. aeruginosa* translational activity in an amino acid-rich growth environment.

**Fig. 1.**
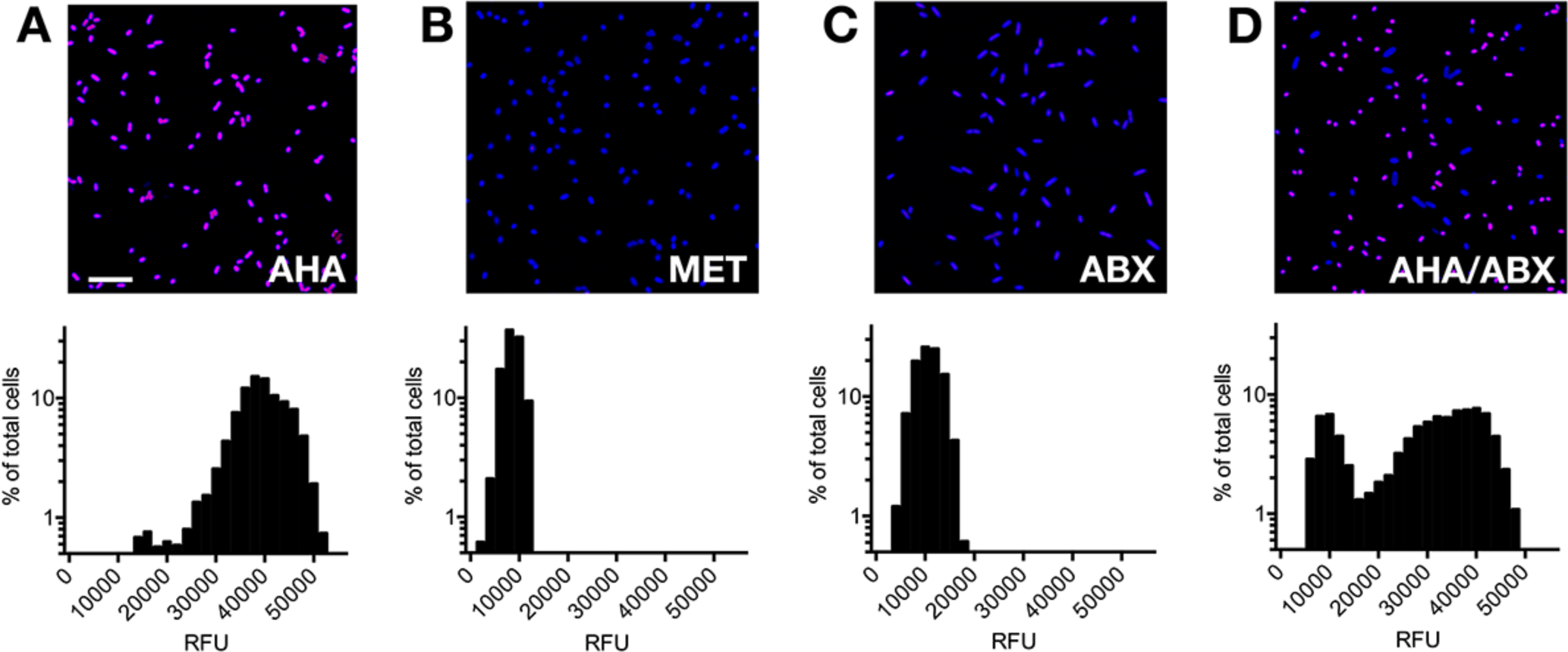
BONCAT labeling of *P. aeruginosa* differentiates translationally active and inactive cells. *P. aeruginosa* was incubated in the presence of (**a)** AHA, (**b)** methionine (MET), and (**c)** antibiotics prior to AHA (ABX). Actively growing cells were identified via strain-promoted ‘click’ chemistry (Cy5, magenta; SYTO64, blue). Histograms associated with each image represent average Cy5 pixel intensity (relative fluorescence units, RFU) per cell. (**d)** Two AHA-treated cultures (one with antibiotics, one without) were mixed in a 1:1 ratio prior to Cy5-DBCO labeling. These data demonstrate that BONCAT can differentiate translationally active and inactive bacterial cells in a complex nutritional milieu. Scale bar = 10µm.

To assess whether BONCAT is broadly suitable for labeling polymicrobial communities found in the CF airways, we then performed mixed activity labeling as described above on representative isolates of *Achromobacter xylosoxidans, Burkholderia cenocepacia, Escherichia coli, Fusobacterium nucleatum, Prevotella melaninogenica, Rothia mucilaginosa, Staphylococcus aureus, Stenotrophomonas maltophilia, Streptococcus parasanguinis*, and *Veillonella parvula* (Fig. 2). Each mixed culture (+/- antibiotics in a 1:1 ratio) exhibited a similar labeling pattern to *P. aeruginosa*, suggesting that BONCAT can be used to characterize translational activity among diverse bacterial taxa associated with the CF airways. Importantly, BONCAT labeling did not affect the growth phenotype of any species under our experimental conditions (Supplementary Fig. 3), consistent with previous studies showing that BONCAT permits labeling of microbiota without concomitant changes in growth rate or protein expression (36,44).

**Fig. 2.**
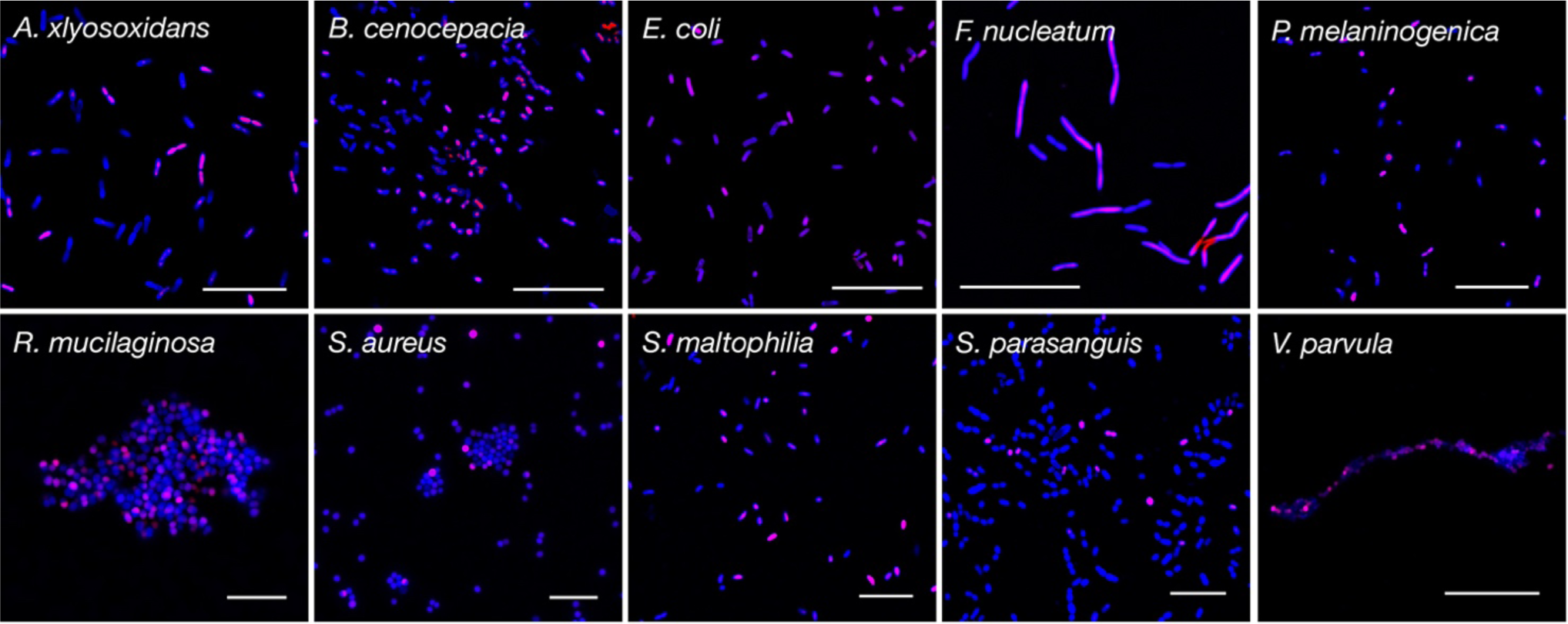
BONCAT can identify active cells among diverse CF microbiota. Two cultures (one treated with antibiotics, one without) of each species were grown in the presence of AHA and mixed 1:1 prior to Cy5-DBCO (magenta) labeling and SYTO64 counterstaining (blue). These data demonstrate that BONCAT can differentiate between active and inactive bacterial cells among diverse CF microbiota. Scale bars; *Ax, Bc, Fn, Ec, Pm, Rm* = 20µm; *Sa, Sm, Sp =* 10µm; *Vp =* 5µm.

### BONCAT identification of active CF microbiota

The BONCAT protocol developed for lab-grown cultures was then modified for analysis of CF bacterial communities *in situ*. To do so, sputum was collected and immediately supplemented with cycloheximide to reduce AHA incorporation by host cells. Samples were then divided into three equal-volume aliquots, supplemented with either AHA, methionine, or antibiotics (chloramphenicol/tetracycline/tobramycin) plus AHA, and incubated at 37°C for 3h. Incubation time was chosen to maximize labeling while minimizing changes in bacterial growth conditions such that they closely reflected the *in vivo* chemical environment. AHA concentration (6mM) was based on average methionine content in CF sputum (0.6mM)(45) and a 10:1 AHA:MET ratio (or greater) required for effective labeling (Supplementary Fig. 4).

Representative micrographs (Fig. 3a) reveal BONCAT labeling of bacterial cells within sputum obtained from three individual CF subjects (Supplementary Table 1, Patients 1-3). Consistent with previous reports of heterogeneous growth rates *in vivo* (25,26), notable differences in Cy5 fluorescence are apparent at higher magnification (Fig. 3b); several individual cells and cell aggregates show moderate to intense labeling whereas others are unlabeled. Treatment with methionine instead of AHA did not result in fluorescent signal, ruling out non-specific labeling (Supplementary Fig. 5). Similarly, treatment of sputum with antibiotics prior to AHA addition also resulted in a significant reduction in fluorescence intensity. However, this reduction was incomplete, which may reflect the development of antimicrobial tolerance that arises among CF pathogens over time (Supplementary Fig. 5). Average pixel intensity per bacterial cell (Fig. 3c) further emphasizes the range of bacterial translational activity *in situ* and the likely slower growth rates of CF microbiota compared to cultures grown *in vitro* (compare Fig. 3c and Fig. 1a). These analyses demonstrate that BONCAT labeling can be used to characterize bacterial activity within complex sputum samples. Moreover, these data suggest that translationally-active bacteria represent only a subpopulation of the CF lung microbiota and exhibit heterogeneous growth activity *in situ*.

**Fig. 3.**
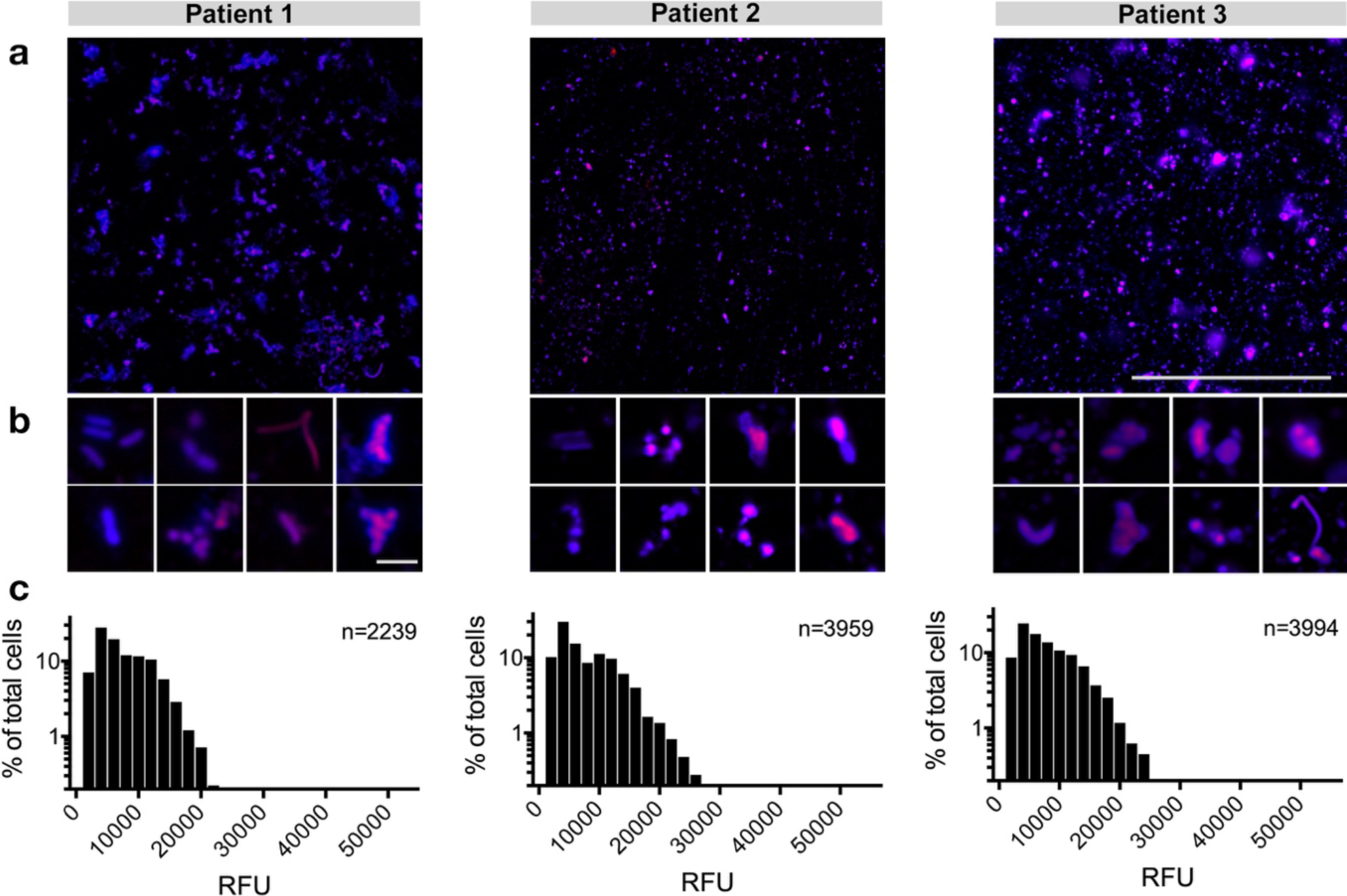
CF microbiota exhibit heterogeneous translational activity within sputum. (**a**) Sputum was incubated in the presence of 6mM AHA immediately upon expectoration. BONCAT labeling with Cy5-DBCO (magenta) and counterstaining with SYTO64 (blue) reveals heterogeneous AHA incorporation (*i.e.* translational activity) by CF microbiota. (**b**) Higher magnification images further emphasize the range of bacterial activity at the single-cell level. (**c**) Average Cy5 pixel intensity per cell suggests slow and heterogeneous growth rates of bacterial cells *in situ*. Scale bars; a = 100µm, b = 5µm.

### Flow cytometric analysis of BONCAT labeled CF microbiota

BONCAT combined with fluorescence-activated cell sorting (FACS) has previously been used to study microbial activity within soils and marine sediments (35,38). We therefore sought to use FACS to characterize and isolate BONCAT labeled (*i.e.* active) cells within sputum samples derived from clinically stable CF subjects at various stages of disease(Supplementary Table 1, patients 4-7). Our experimental workflow is shown in Figure 4. Upon sputum collection, a small aliquot (“original”) was removed and stored at −80°C prior to conventional 16S rRNA amplicon analysis. The remaining sample was then supplemented with AHA for 3h, subjected to Cy5-DBCO labeling and counterstained, followed by removal of another aliquot (“sort input”) that was used to determine community profile changes as a result of sputum incubation *ex vivo*. The remaining sample was then homogenized and filtered to remove host cells, followed by FACS to isolate Cy5-(“sort negative”) and Cy5+ (“sort positive”) cells. Sort integrity was validated by immunostaining using an anti-Cy5 antibody to identify Cy5+ (*i.e.* active) cells (Supplemental Fig. 6).

**Fig. 4.**
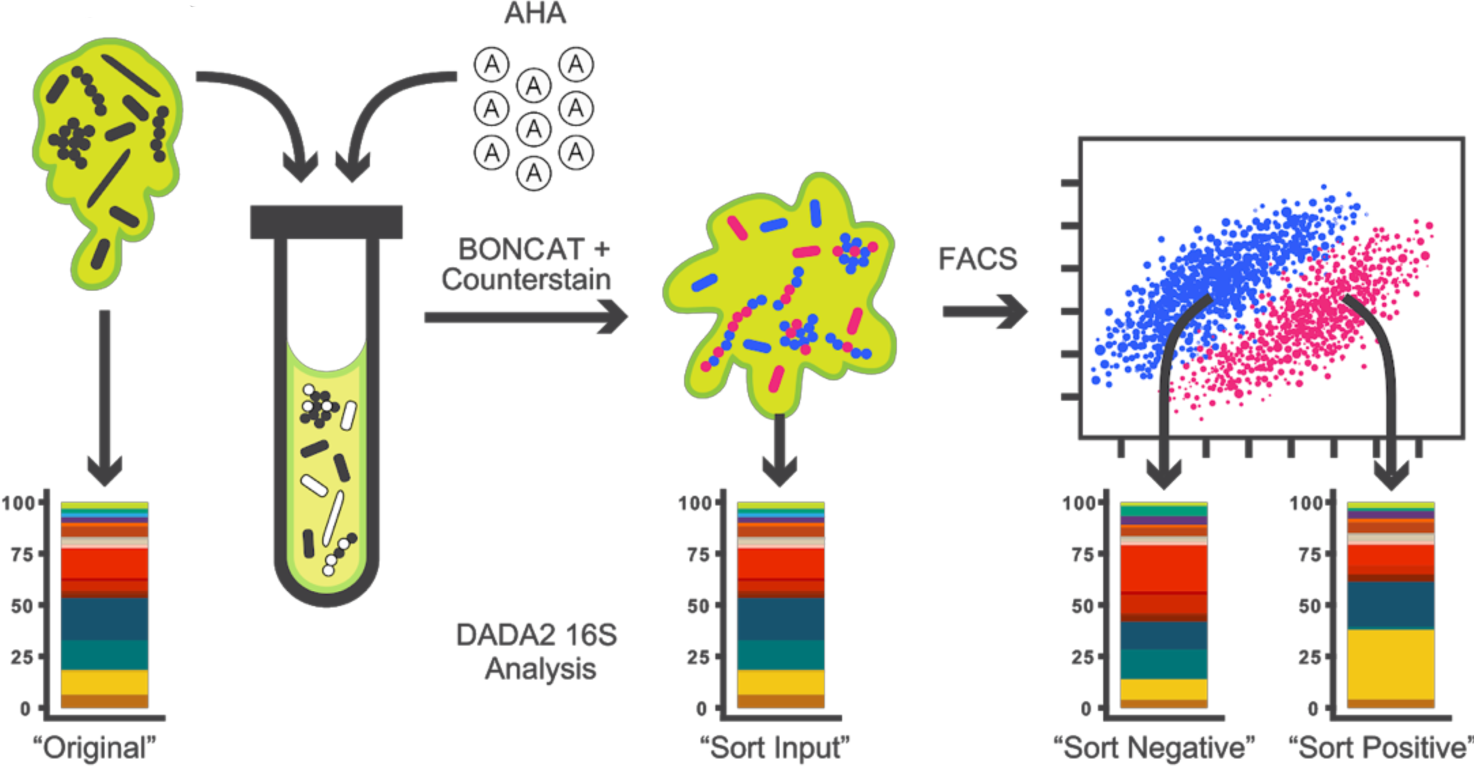
Experimental workflow for BONCAT analysis of CF sputum.

BONCAT-FACS plots of CF sputum samples are shown in Figure 5a. Cy5-and Cy5+ gates were patient-specific and were established first by using an AHA-aliquot to define the negative gate for each sample. Positive gates were then conservatively assigned by comparing the AHA+ sample to the AHA-control (see Supplementary Fig. 1 for gating scheme). The AHA+ sample underwent a notable shift along the Cy5+ axis, reflective of translational activity (Fig. 5a). Cells that fell in the Cy5+ gate exhibited a higher geometric mean of fluorescence intensity in the Cy5 channel (relative to the Cy5-gate), confirming BONCAT labeling (Supplementary Fig. 8). Based on fluorescent events, we generally found that only a small subset of the overall bacterial population was Cy5+ (12.1-18.5% of the parent population; Supplementary Table 2). These data reflect labeling patterns shown by microscopy (Fig. 3) and suggest that expectorated sputum harbors only a small percentage of translationally active cells.

**Fig. 5.**
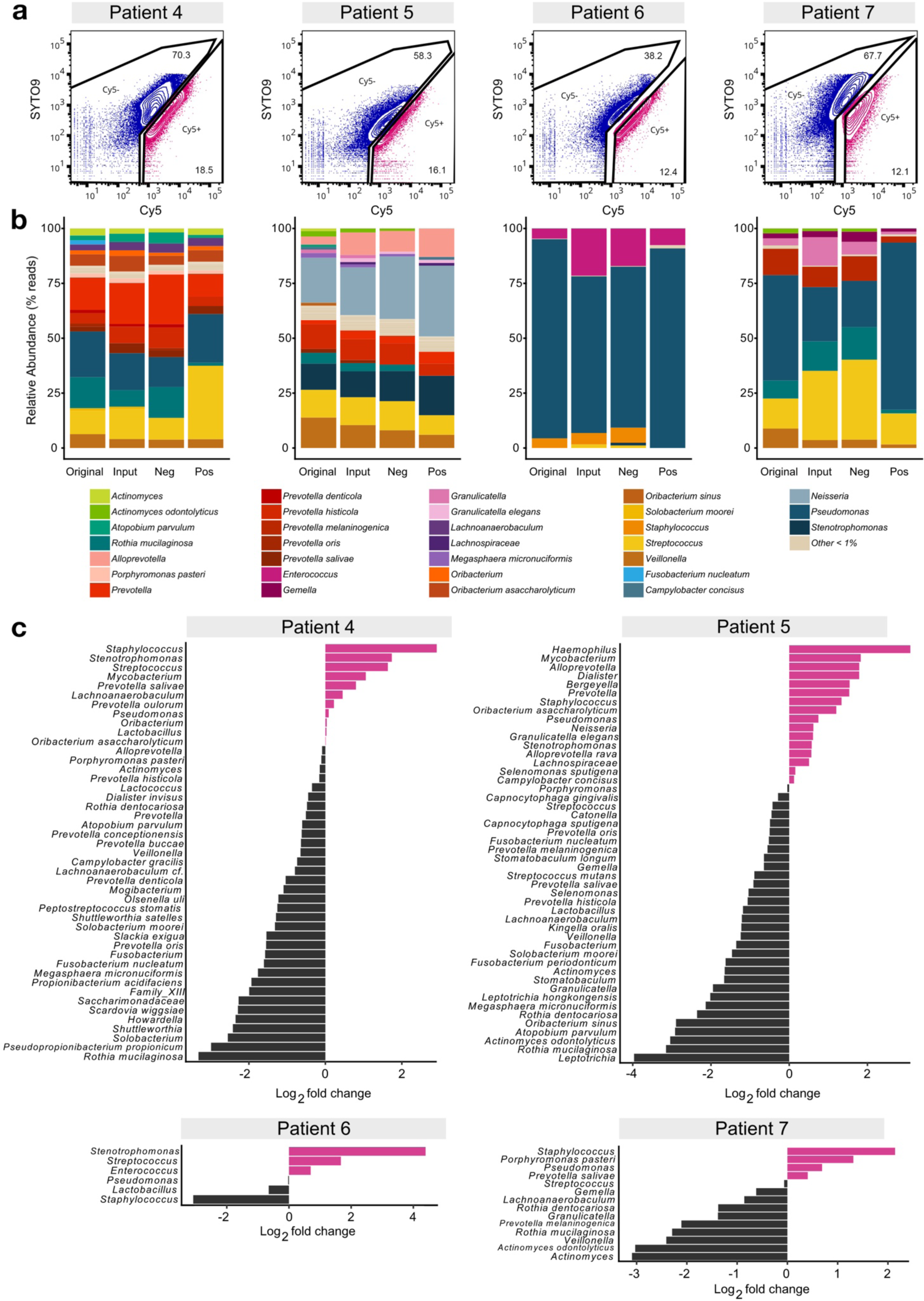
BONCAT labeling, FACS and 16S rRNA sequencing of CF sputum reveals the taxonomic identities of metabolically active microbiota. **(a)** FACS of BONCAT labeled sputum reveals Cy5-and Cy5+ subpopulations. Percentages shown reflect % of parent population post-CD45RO gating. **(b)** The “original”, “sort input”, “sort negative”(Cy5-), and “sort positive”(Cy5+) fractions were analyzed by 16S rRNA gene sequencing and the DADA2 algorithm. Taxa plots summarize sequencing data by patient and sorted fractions. **(c)** Log2 fold changes between relative abundances of taxa in the “sort positive” compared to the “original” fraction. Pink bars indicate taxa that were increased in relative abundance in the “sort positive” fraction, representing translationally active microbiota.

### 16S rRNA gene sequencing reveals the taxonomic identities of active CF sputum microbiota

To determine bacterial community composition, 16S rRNA gene sequencing was applied to “original”, “sort input”, “sort negative” and “sort positive” fractions from each sample. Sequence data were analyzed using the DADA2 pipeline (46) to resolve amplicon sequence variants instead of the more common approach of clustering amplicon sequences into operational taxonomic units (OTUs). DADA2 reduces potential loss of meaningful biological sequence variation due to clustering by similarity and can improve the ability to observe fine-scale variation (including species-level resolution) in bacterial populations. Using this approach, sequence data derived from AHA labeled samples (“sort input”, “sort negative”, “sort positive”) were compared to their paired “original” sample to characterize the translationally active subpopulations.

Each subject harbored lung microbiota of moderate complexity (Fig. 5b), and community profiles were consistent with prior 16S rRNA gene surveys of CF sputum in which *Pseudomonas, Streptococcus*, and *Staphylococcus* were dominant genera (3-5,7-12). We also achieved species-level resolution for less abundant taxa, including several obligate and facultative anaerobes (*e.g. Prevotella sp., F. nucleatum*). In general, AHA-labeling did not result in substantial changes in bacterial membership; for the most abundant taxa (>5%), community composition was comparable before (“original”) and after (“sort input”) BONCAT labeling, demonstrating that AHA treatment for 3h had minimal effect on relative bacterial abundance. Interestingly, bacterial populations recovered from FACS analysis (“sort negative”, “sort positive”) also showed strong similarity among the most abundant community members relative to the original sample (*i.e.* those detected by conventional sequencing). Less abundant taxa (1-5%, Supplementary Fig. 7), showed greater variation between fractions, but most were also generally detectable in both sort negative and sort positive gates. Together, these data suggest a subset of most taxa detected by conventional 16S rRNA sequencing are translationally active *in vivo*. Moreover, each taxon appears to exhibit heterogeneous growth activity that may have important implications for disease progression and patient therapy.

To better observe changes in the relative abundance of bacterial taxa between fractions, we calculated the log2 fold change (log2fc) between “original” and “sort positive” as well as differences between “sort positive” and “sort negative” for each patient (Fig. 5c, Supplementary Fig. 9). Some genera/species present in the log2fc plots do not appear in taxa plots (Fig. 5b) because they were less than 1%, but we note that activity among less abundant populations may also be a determinant of CF pathogenesis. In Patient 4 for example, *Stenotrophomonas* and *Mycobacterium sp.* together make up less than 0.04% of the overall bacterial community but were enriched ∼1.7 and 1.1 log_2_fold in the active fraction, respectively. *Streptococcus* sp. was one of the more abundant genera in the original community (10.8%) and represented an even greater percentage of the active population (33%, or 1.6-log_2_fold enrichment). Conversely, *R. mucilaginosa*, which was also abundant (14.4%), showed little translational activity as determined by BONCAT labeling (as it did in patients 5 and 7).

Patients 5-7 exhibited other notable differences in bacterial abundance between fractions (Fig. 5c, Supplementary Fig. 9). In Patient 5, *Haemophilus* and *Mycobacterium* showed the greatest enrichment in the active fraction (3.2 and 1.8-log_2_fold enrichment, respectively) but again only made up a small percentage of the original community (0.1%). *R. mucilaginosa* and *Leptotrichia*, present at 5.2% and 0.1% respectively, were at much lower abundance (3.1 and 4.0-log_2_fold reduction) in the active fraction. Patient 6 harbored a *Pseudomonas*-dominated (90.6%) community in which *S. aureus* was also abundant (4.4%), but the latter exhibited little translational activity (3.1-log_2_fold decrease) suggesting a predominantly dormant *S. aureus* population. *Stenotrophomonas* and *Streptococcus* exhibited the greatest enrichment (1.7 and 4.4-log_2_fold) in the active fraction of Patient 6, despite being present at low abundance. Finally, Patient 7, whose sputum contained a diverse bacterial community composed of 26 identifiable bacterial taxa, harbored an active subpopulation that was dominated by *P. aeruginosa* (74.7%, 0.7-log_2_fold increase) and *Streptococcus* (15.2%, no change).

Taken together, BONCAT data reveal the extensive heterogeneity of translational activity among CF microbiota. Each patient harbors a unique bacterial community, though community membership and relative abundance is not necessarily predictive of active growth. Ultimately, profiling of bacterial communities in this manner may help guide therapeutic strategies by identifying subpopulations of translationally active bacteria.

## DISCUSSION

16S rRNA gene sequencing has become the gold standard for culture-independent characterization of CF airway bacterial communities. Despite the wealth of data that has emerged regarding the complexity of lung microbiota, we have little understanding of *in vivo* bacterial activity and the specific contributions of individual species to pathogenesis. Expanding on recent studies employing BONCAT as a means of characterizing the ecophysiology of microbial communities in their natural growth environment (35-38), we use this approach in combination with FACS and 16S rRNA gene sequencing to shed light on bacterial activity in CF sputum. We demonstrate that only a subset of each taxon is detectable by metabolic labeling. Identification and characterization of this subpopulation is not achievable using conventional sequencing approaches and may provide a more precise representation of relevant microbiota within the CF lung.

BONCAT-based studies of translationally active bacteria challenge our thinking on the microbial ecology of the CF airways. Each subject harbored a unique bacterial community consisting of canonical lung pathogens (e.g. *Pseudomonas, Staphylococcus, Achromobacter, Mycobacterium, Streptococcus*). Consistent with previous studies using RNA-based methods (24,25), BONCAT-FACS-based sequencing data indicate these most abundant taxa are also active *in situ*, reinforcing their probable role in CF pathogenesis. However, we also revealed low abundance community members comprise a disproportionate percentage of the “sort positive” fraction despite a quantitatively low abundance in the overall (“original”) bacterial community. In conventional 16S rRNA datasets, ‘rare’ taxa (*i.e.* <1%) can be challenging to detect among high abundance organisms (or they are grouped into an ‘other’ category). Moreover, given the abundance of dead and/or dormant cells observed in this study (∼38-70%; Supplementary Table 2), longitudinal dynamics among low abundance taxa would be masked in standard taxa plots, and may partially explain why observed within-patient differences in bacterial community composition rarely track with patient symptoms (9,15-22). We hypothesize that low abundance organisms represent keystone members of the lung microbiota, whose *in vivo* activity dynamics are determinants of acute inflammation, either by directly impacting the host, or indirectly through modulating the growth and virulence of higher abundance pathogens.

BONCAT imaging of sputum and log2fc plots between positive and negative gates demonstrated both population-wide and taxon-specific translational heterogeneity. This spectrum of translational activity may confer a significant advantage for bacteria and optimize their fitness in the complex environment of the CF lung. Airway microbiota face a dynamic milieu shaped by microbial competitors, antimicrobials, the host immune response, nutrient limitation and other chemical stimuli that can be unfavorable to growth. Under these conditions, adopting a ‘bet hedging’ strategy in which only a subpopulation of cells is active may ensure that a given bacterial species is prepared to contend with environmental stress (47). In addition, the transition between translationally active and dormant states may help explain the periodicity of PEx; faced with a favorable growth environment, more cells of a given taxon (or taxa) may be induced into active growth and elicit a heightened patient response.

The balance between growth states may also be a critical determinant of patient response to therapy. By adopting a ‘persister’-like strategy in which reduced cellular activity confers a temporary multidrug-resistant phenotype, a dormant subpopulation could ensure persistence during an antibiotic challenge. Once antibiotic selective pressure is relieved, antimicrobial tolerant populations may emerge. This heterogeneity may also help explain instances where a patient’s clinical response is not predicted by the *in vitro* drug susceptibility of a given pathogen. We posit that clinical sensitivity panels are poorly predictive of antibiotic efficacy because they do not account for the heterogeneous *in situ* translational activity described here.

While active cells are likely more responsible for pathogenesis, inactive cells (Cy5-) are also of importance to CF lung disease as bacteria do not necessarily have to be translationally active to influence their greater community. For example, it is known that largely dormant populations can drive geochemical processes in their growth environment (*e.g.* mineralizing organic C to CO_2_)(48). Translationally inactive cells can also shape their growth environment through nutrient exchange, secretion of virulence factors and small metabolites, electrostatic interactions, and stimulation of the host immune response. Further characterization of *in situ* activity heterogeneity, the contributions of both active and dormant populations to disease, the frequency of transition between states and the factors that stimulate those transitions will help us better understand disease dynamics.

Though BONCAT represents a useful tool for the study of CF microbiota, there are notable limitations. First, bacterial cell sorting by flow cytometry is imperfect, as each species has characteristic sort properties. When defining our gating scheme, Cy5+ and Cy5-gates were conservatively selected (requiring a gap in between gates) such that the selection of “inactive” cells in the positive gate is minimized, and vice versa. However, with this gap a subset of the active population is not collected. Similarly, there is a high probability of selecting “active” cells in the negative gate due to flow migration characteristics (*e.g. F. nucleatum* shifts differently than a much smaller *V. parvula* cell). Finally, bacterial aggregates, in which only some cells are active (see Fig. 3b) could be pulled into the negative gate by the inactive population of that aggregate. We are currently exploring alternative approaches to improve upon the sorting efficiency of BONCAT labeled cells.

It is also possible BONCAT is selective against certain taxa. While sputum chemistry is stable over time *ex vivo* (43), it is notable that after AHA incubation, patients 4, 5, and 7 had a decrease in abundance of facultative and obligately anaerobic taxa (*e.g. Fusobacterium, Veillonella* and *Rothia*) in “sort positive” fractions relative to the original sample (Fig. 5c). However, this was not always the case (*e.g.* some species of *Prevotella* and *Porphyromonas* increased) making it difficult to determine whether the observed log2 fold changes reflect growth constraints during BONCAT labeling or a true slow growth (or dormant) phenotype. Though each bacterium tested *in vitro* demonstrated the ability to uptake AHA (Fig. 2), it is expected that each species will incorporate AHA into new proteins at different rates. Future work will be aimed at optimizing reaction conditions and incubation times to minimize the effect of the experimental approach biasing FACS and sequencing data.

Despite these limitations, BONCAT can be used to extend our understanding of the role of specific microbiota in CF lung disease. Here we focused on a cross-sectional cohort of stable subjects, but the approach can be used to address important questions about microbial community dynamics over time. For example, (1) how do active populations vary with disease state? Future studies will focus on longitudinal analyses of within-subject microbial dynamics and how active species correlate with patient symptoms. By identifying bacterial subpopulations most active either preceding or during an acute disease flare (*i.e.* PEx), more effective therapeutic strategies are likely to be identified. (2) Why are only some patients responsive to antimicrobial therapy? As mentioned above, *in vivo* drug efficacy is often inconsistent with clinical sensitivity panels. By obtaining sputum and amending small aliquots with different classes of antibiotics, BONCAT analysis of the ensuing changes in bacterial activity can be used to predict how CF subjects might respond to treatment. (3) How do specific taxa respond to environmental stimuli? It is known that bacteria are dynamically responsive to their growth environment, yet how CF microbiota adapt to perturbations in the sputum milieu is poorly understood. BONCAT characterization of sputum samples amended with specific nutrients or incubation under varying environmental conditions (*e.g.* low pH) will help shed light on parameters that constrain or potentiate bacterial growth *in vivo*. (4) How is translational activity spatially arranged? With the exception of small bacterial aggregates (Fig. 3), the approach described here offers limited insight on the spatial distribution of bacterial activity. As an alternative to FACS-based sequencing, BONCAT could be combined with fluorescent in situ hybridization (FISH) methods and histological analysis of sputum (or lung tissue) to visualize spatial relationships between actively growing bacteria.

In summary, we demonstrate that BONCAT is a powerful tool for the visualization and identification of translationally active bacteria and provides a measure of microbial activity not captured by conventional molecular profiling. Our use of BONCAT lays the foundation for a more detailed understanding of the *in situ* physiology of CF microbiota and has important implications for the development of new therapeutic strategies and improved clinical outcomes. In addition, the approach is broadly applicable to other airway diseases (*e.g.* COPD, ventilator associated pneumonias, and sinusitis) where the activity of complex bacterial communities is central to disease states. We are currently using this approach to study microbial community dynamics in a variety of infectious disease contexts.

## MATERIALS AND METHODS

### Bacterial strains and culture conditions

Bacterial strains are listed in Table 1. *Fusobacterium nucleatum, Prevotella melaninogenica, Veillonella parvula*, and *Streptococcus parasanguinis* were derived from the American Tissue Type Collection and obtained from Microbiologics (St. Cloud, MN). *Rothia mucilaginosa* was obtained from the Japan Collection of Microorganisms (Riken, Tokyo). *Staphylococcus aureus, Escherichia coli* and *Pseudomonas aeruginosa* were obtained from D.K. Newman (California Institution of Technology), and *Burkholderia cenocepacia* was obtained from C.H. Mohr (University of Minnesota). *Achromobacter xylosoxidans* and *Stenotrophomonas maltophilia* were isolated from patients undergoing treatment at the UMN Adult CF Center. Aerobes were maintained on Luria-Bertani (LB) agar, while anaerobes were maintained on Brain-Heart Infusion (BHI) agar supplemented with a 5% vitamin K-hemin solution (Hardy Diagnostics #Z237) in an anaerobic chamber (Coy) under a 90% N_2_ / 5% CO_2_ / 5% H_2_ atmosphere.

**Table 1.**
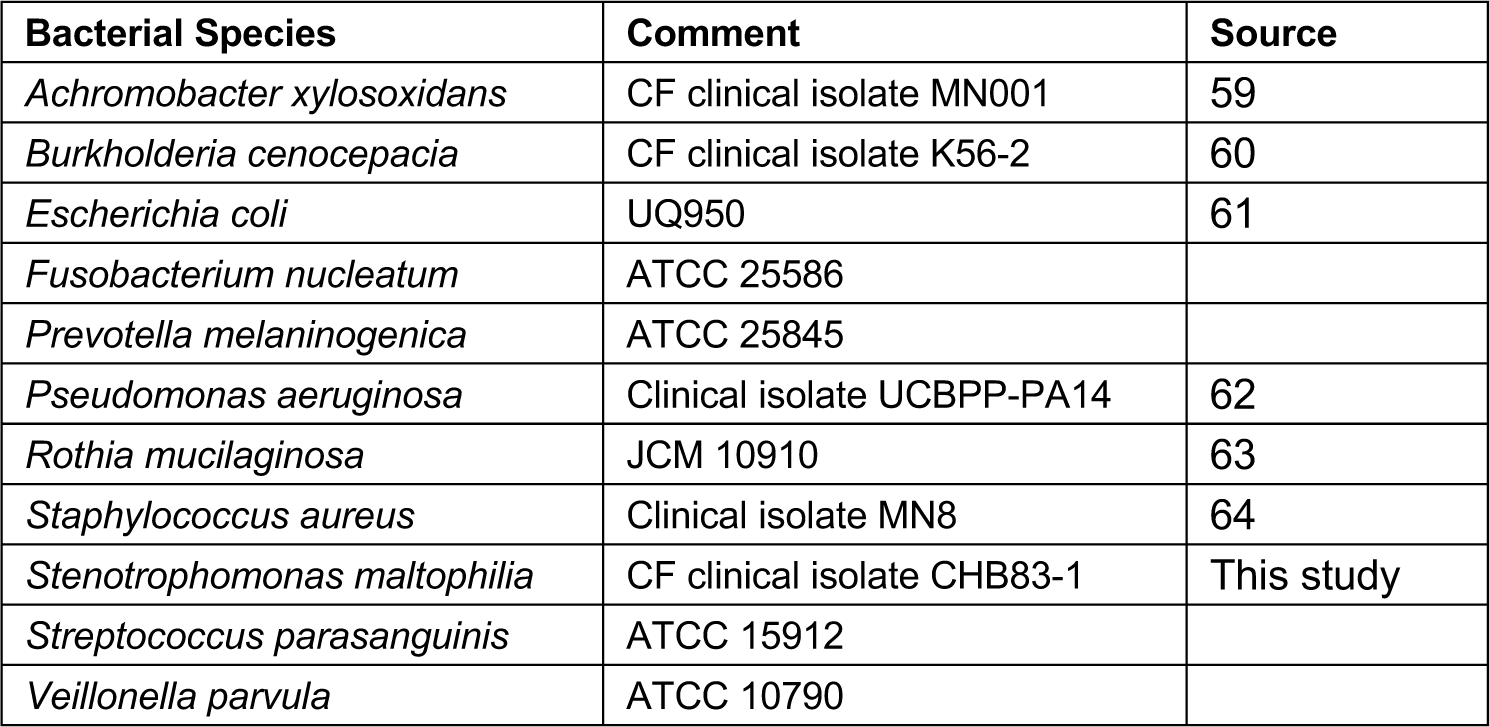
Bacterial strains used in this study.

### Clinical sample collection

Spontaneously expectorated sputum was collected from stable subjects with cystic fibrosis during routine outpatient visits to the Adult CF Center at the University of Minnesota. All subjects provided written informed consent prior to sample collection as approved by the UMN Institutional Review Board (Study #1403M49021). Upon consent, each patient provided a single sample that was collected in a sterile 50mL conical tube. Patient data are shown in Supplementary Table 1.

### Bioorthogonal non-canonical amino acid tagging (BONCAT)

BONCAT labeling was performed as described by Hatzenpichler (35,36) with modifications. For imaging of lab-grown cultures (see below), *P. aeruginosa, B. cenocepacia, A. xylosoxidans, S. maltophilia, R. mucilaginosa, E. coli and S. aureus* were grown aerobically in LB, while *S. parasanguinis, V. parvula, P. melaninogenica* and *F. nucleatum* were cultured under anaerobic conditions in BHI broth supplemented with hemin and vitamin K. Cultures were grown overnight and diluted 1/100 in 10mL of fresh medium. Upon reaching mid-log phase, cultures were supplemented with either 6mM AHA or 6mM methionine (MET) and incubated for 3h at 37°C. When indicated, an antibiotic cocktail consisting of chloramphenicol (30µg/mL), tetracycline (200µg/mL) and tobramycin (10µg/mL) was added 30 minutes prior to AHA addition to arrest protein synthesis. After incubation, cultures were pelleted via centrifugation (5 min at 10,000 × *g*), fixed in 4% paraformaldehyde (PFA) for 2h at 4°C, resuspended in phosphate buffered saline (PBS, pH 7.4) and stored at 4°C.

Patient sputum samples used for imaging were treated with cycloheximide (100µg/mL) upon expectoration and divided into three equal volumes. Aliquots were supplemented with either AHA (6mM), methionine (6mM), or AHA (6mM) with chloramphenicol/tetracycline/tobramycin as described above, incubated at 37°C for 3h, followed by fixation in 4% PFA overnight at 4°C. Samples collected for flow cytometry were divided into three 500µL aliquots. One control aliquot was immediately frozen at −80°C and later used for conventional 16S rRNA gene sequencing. Cycloheximide (100µg/mL) was added to the remaining two aliquots, one of which was supplemented with AHA (6mM) and incubated at 37°C for 3h. Labeled samples (and unlabeled controls) were then fixed in 4% PFA for 2h, pelleted via centrifugation (5 min at 10,000 × *g*), resuspended in PBS, stored at 4°C and further processed within 24h.

### Click chemistry

For each bacterial culture and sputum sample, strain-promoted azide-alkyne cycloaddition (‘click’ chemistry) (49) was also performed as described previously (36). Briefly, fixed biomass was pelleted, resuspended in freshly prepared 2-chloroacetamide (100mM) and incubated for 1h at 46°C, shaking at 450rpm in the dark. Cy5-dibenzocyclooctyne (Cy5-DBCO) (Click Chemistry Tools) was then added to a final concentration of 10µM followed by incubation for 30 min at 46°C. Samples were washed three times in PBS and further processed for imaging and flow cytometry (see below).

### SDS-PAGE

*P. aeruginosa* was grown to late-exponential phase as described above and supplemented with varying concentrations of AHA (100µM to 1mM) for 1h prior to fixation. Similarly, *P. aeruginosa* was grown in the presence of varying ratios of MET:AHA. Bacterial pellets were resuspended in extraction buffer (1% sodium dodecyl sulfate, 50mM NaCl, 100mM EDTA, 1mM MgCl_2_ at pH 8.4) and boiled for 30 min. After boiling, samples underwent click chemistry as described above. A mixture of methanol:chloroform:water (12:3:8) was then added to each sample followed immediately by centrifugation for 5 min at 16,000 × g. The water/methanol phase was then carefully removed, and protein recovered from the interface was washed 3 times in 100% methanol. After the final wash, supernatant was removed and pellets were air dried. Protein was resuspended in 100μl 1X LDS (lithium dodecyl sulfate) sample buffer and denatured at 70°C for 10 minutes. 10μl of protein was run on an 8% Bis-Tris gel with MOPS (3-(*N*-morpholino)propanesulfonic acid)-sodium dodecyl sulfate (SDS) running buffer to which sodium bisulfite had been freshly added. Gels were run at 150V, fixed for 30 min in a 1:2:7 acetate:methanol:water mix, and imaged with a Typhoon FLA 9500 scanner (GE Healthcare) using an excitation wavelength of 635nm.

### Fluorescence microscopy

BONCAT labeled bacterial cultures and sputum were spotted on Superfrost Plus microscope slides and counterstained using 1.6µM STYO64 in PBS. Slides were then washed twice in PBS, mounted using Prolong Diamond Antifade and imaged using an Olympus IX83 microscope with a transmitted Koehler illuminator and a 60X oil objective lens (NA 1.42). Images were captured on a Hamamatsu ORCA-Flash4.0 V2 digital CMOS camera, and post-acquisition image analysis was performed using cellSens software (v.1.14, Olympus). SYTO64 and Cy5 were visualized using excitation/emission wavelengths of 562nm/583nm and 628/640nm, respectively.

Image analysis was performed using FIJI (50). Briefly, images were subjected to background subtraction using a rolling ball radius of 150 pixels. Individual cells were identified by adjusting thresholds of SYTO64 images using Huang’s fuzzy thresholding method (51). Images were also segmented using a watershedding algorithm that assumes each maximum belongs to a discrete particle. The ‘*Analyze Particles’* operation was used to detect and record locations of individual bacterial cells in a given image. For clinical samples, particles were constrained between 100-1000 pixels to minimize detection of host cells and sputum debris. Mean pixel intensity at 647nm (Cy5) was then quantified for each assigned particle. Imaging experiments were performed in triplicate for each bacterial species, and twenty images for each sample were captured (n>2700 particles per sample).

### Flow Cytometry

Prior to sorting, Cy5-DBCO labeled sputum was collected by centrifugation and counterstained with 1.6µM SYTO9 (Invitrogen) in PBS for 30 min. Sputum samples were also stained with 1 µg/ml of PE anti-human CD45RO in PBS (BioLegend) for 30 min to stain activated and memory T cells, some B cell subsets, activated monocytes/macrophages, and granulocytes. All samples were washed in PBS, homogenized using 16-and 22-gauge needles and filtered through a 40μm cell strainer. To separate AHA+ and AHA-bacterial populations, clinical samples were analyzed and sorted on a FACSAriaIIu Cell Sorter (Beckton Dickinson) with a 70μm nozzle at 70psi. Contaminating human leukocytes staining positive for PE anti-human CD45RO were excluded from bacterial populations of interest in the initial sorting gate (Supplementary Fig. 1). An AHA-control was then matched to each sample to determine the level of non-specific Cy5-DBCO binding and was used to establish Cy5+ (*i.e.* active) and Cy5-(*i.e.* inactive) sorting gates. Forward scatter and side scatter gates were then applied to remove large particulates and debris, and liberal doublet discrimination was used to minimize loss of bacterial aggregates. Collected samples were stored at 4°C and processed within 24h. FlowJo software (v.10.5.0) was used for data analysis and presentation.

Cy5+ and Cy5-sorted populations were assessed for post-sort purity by flow cytometry, while collected fractions were visualized by anti-Cy5 immunostaining. To do so, BONCAT labeled sputum samples were spread across Superfrost Plus microscope slides using a sterile pipette tip and allowed to air dry for 30 min. Slides were washed 3X in PBS and blocked using 1% goat serum in PBS for 1h, followed by treatment with an anti-Cy5 monoclonal antibody (C1117, Sigma-Aldrich)(1:100 dilution) in incubation buffer (1% goat serum, 0.3% Triton X100 and 10mg/mL bovine serum albumin) overnight at 4°C. Slides were washed 3X, and incubated with Alexa Fluor 488 goat anti-mouse secondary antibody (1:250) in incubation buffer for 45 min. Slides were washed 2X, counterstained using 0.1% DAPI in PBS and mounted using Prolong Diamond Antifade. Slides were imaged as described above.

### DNA extraction

Genomic DNA (gDNA) was extracted using a modified phenol-chloroform method previously described (52). Briefly, FACS-sorted samples were collected onto 0.22µm polycarbonate membranes (EMD Millipore), which were then transferred to 1 mL of TENS buffer (50mM Tris-HCl [pH 8.0], 20mM EDTA, 100mM NaCl, 1% SDS) containing lysozyme (0.2mg/mL) and lysostaphin (0.02µg/mL) and incubated at 37°C for 30 min. Sodium dodecyl sulfate (SDS) and proteinase K were added to final concentrations of 1% and 1.2mg/mL, respectively, and samples were incubated overnight at 55°C. Enzymes were deactivated by incubating samples at 90°C for 30 min, and sample liquid (including membrane) was transferred to a 5mL conical tube containing an equal volume of phenol:chloroform:isoamyl alcohol (P:C:I, 25:24:1, pH 7.9), which dissolved the membrane. The resulting sample was then split into two Lysing Matrix E tubes (MP Biomedicals) and processed twice by bead beating for 30 seconds. Contents of both tubes were recombined and centrifuged at 3,200 × g for 20 min. The aqueous layer was transferred to a new tube and P:C:I extraction was repeated, followed by a chloroform:isoamyl alcohol (24:1) extraction. A 1/10th volume of sodium acetate (3M, pH 5.2) was then added and nucleic acid was precipitated using one volume of isopropanol followed by centrifugation at 21,130 x g for 20 min. Supernatant was removed, the pellet was washed with 80% ethanol, and centrifuged at 21,130 x g for 10 min. Finally, the gDNA pellet was air dried, resuspended in 10mM Tris buffer (pH 8.0), and stored at −80°C until sequencing.

### DNA sequencing and analysis

gDNA derived from sputum samples was submitted to the University of Minnesota Genomics Center (UMGC) for 16S rRNA gene library preparation using a two-step PCR protocol described previously (53). The V4 region was amplified and sequenced on an Illumina MiSeq using TruSeq (v.3) 2×300 paired-end technology. FACS sheath fluid and DNA extraction reagent control samples were also submitted for sequencing. These control samples did not pass quality control steps due to DNA content below detection thresholds but were incorporated into downstream analyses. An average of 52,155 sequences per sample were obtained across two sequencing runs. The ‘DADA2’ R package (v.1.2.1) (46) was used to trim and filter sequences, model and correct Illumina sequence errors, align paired-end sequences, and filter chimeric reads. Specifically, the first 20 bases from each sequence were trimmed to remove primer sequences. Forward and reverse sequences were trimmed to 200 bases, maintaining Phred quality scores above 30. All other DADA2 pipeline parameters were run using default options. The resulting amplicon sequence variants (ASVs) were assigned taxonomy using RDP classifier (54) and the SILVA SSU database (Release 132, Dec. 2017)(55,56). The ‘Decontam’ package (v.1.2.0) (57) was used with 16S qPCR data obtained from the UMGC QC protocol to identify sequences that were likely to be processing contaminants. A total of 31 taxa were removed from the dataset based on frequency and prevalence in the sample when compared with DNA extraction and FACS sheath fluid controls. An average of 41,642 sequences per sample were recovered from ‘DADA2’/‘Decontam’ analysis corresponding to 192 unique taxonomic assignments with 76 unambiguously assigned at the species level.

ASV count data and taxonomic assignment were used within the analysis framework of the ‘Phyloseq’ R package (v.1.26.0) (58). ASVs were filtered when they did not belong to the domain Bacteria, or when not assigned taxonomy at the phylum level. Phyla that had low prevalence and abundance (including Verrucomicrobia, Acidobacteria, Deinococcus-Thermus, and Cyanobacteria) were removed from the dataset, as were singleton ASVs. After filtering there remained 105 unique taxonomic assignments with 59 unambiguously assigned at the species level. Log2 fold differences in relative abundance for each taxon were calculated between the “original”, “sort negative” and “sort positive” fractions for each study subject. Taxa that appeared either in the original or positive sort, but not both were excluded from the dataset. To test whether our log2 fold difference analysis was affected by the number of sequences per sample, we also performed this analysis with rarified data and did not observe changes in results. For all figures, a specific epithet was used when assigned exactly from the SILVA database using the lowest possible taxonomic assignment (*i.e.* genus or species).

### Data Availability

Raw 16S rRNA gene sequence data (Fig. 5 and Supplementary Figures 7 and 9) were deposited as fastq files in the NCBI sequence read archive under Bioproject ID PRJNA520921.

## ACKNOWLEDGEMENTS

We thank Roland Hatzenpichler (Montana State University) for technical advice, Alex Vitti for graphic design, the care team at the UMN Adult CF Treatment Center and their patients for participating in the research. This work was supported by a Gilead Sciences Investigator Sponsored Research Award, Cystic Fibrosis Foundation Research Grant (HUNTER16G0) to RCH. KAB was supported by a NIH Lung Sciences T32 fellowship (#2T32HL007741-21) awarded through NHLBI. SKL received support from T32 (#T90 DE0227232) and F31 (#F31 DE027602) fellowships through the National Institute of Dental and Craniofacial Research.

## AUTHOR CONTRIBUTIONS

T.V., S.L., and K.B. contributed equally to this work. T.V., S.L., K.B., and R.H. were responsible for study design and wrote the manuscript. T.V., K.B. and R.H. performed BONCAT labeling and imaging experiments. T.V., K.B., and J.M. performed and optimized flow cytometry and FACS. L.C. and J.D. were responsible for patient recruitment, sample collection and patient data management. S.K. performed sequencing and analysis.

## CORRESPONDING AUTHOR

Correspondence to Ryan C. Hunter.

## Supplemental Data

**Supplementary Figure 1.**
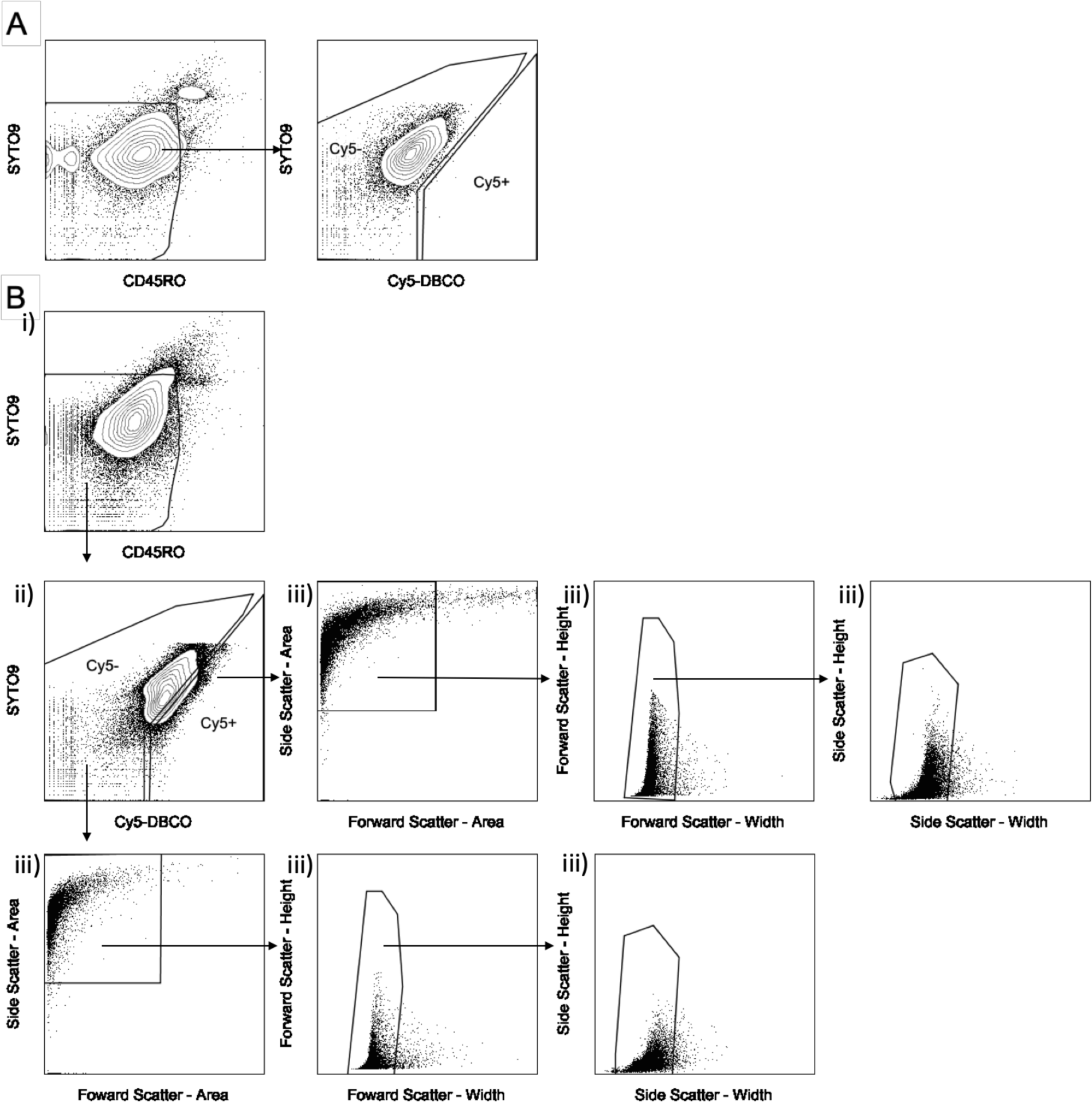
Representative gating scheme for AHA+ and AHA-populations. (a) AHA-control samples were first gated based on PE anti-human CD45RO staining to remove human leukocytes. Next, the negative control was used to measure non-specific Cy5-DBCO binding to define Cy5+ (*i.e.* active) and Cy5-(*i.e.* inactive) sorting gates. (b) shows a patient matched AHA+ sample and the gating used to isolate populations of interest. The gating strategy involved, i) excluding human leukocytes using PE anti-human CD45RO, ii) using Cy5+ and Cy5-gates based on the patient-matched AHA-control, iii) creating forward scatter and side scatter plots to remove large, complex particulates and debris, and liberal doublet discrimination to minimize the loss of bacterial aggregates.

**Supplementary Figure 2.**
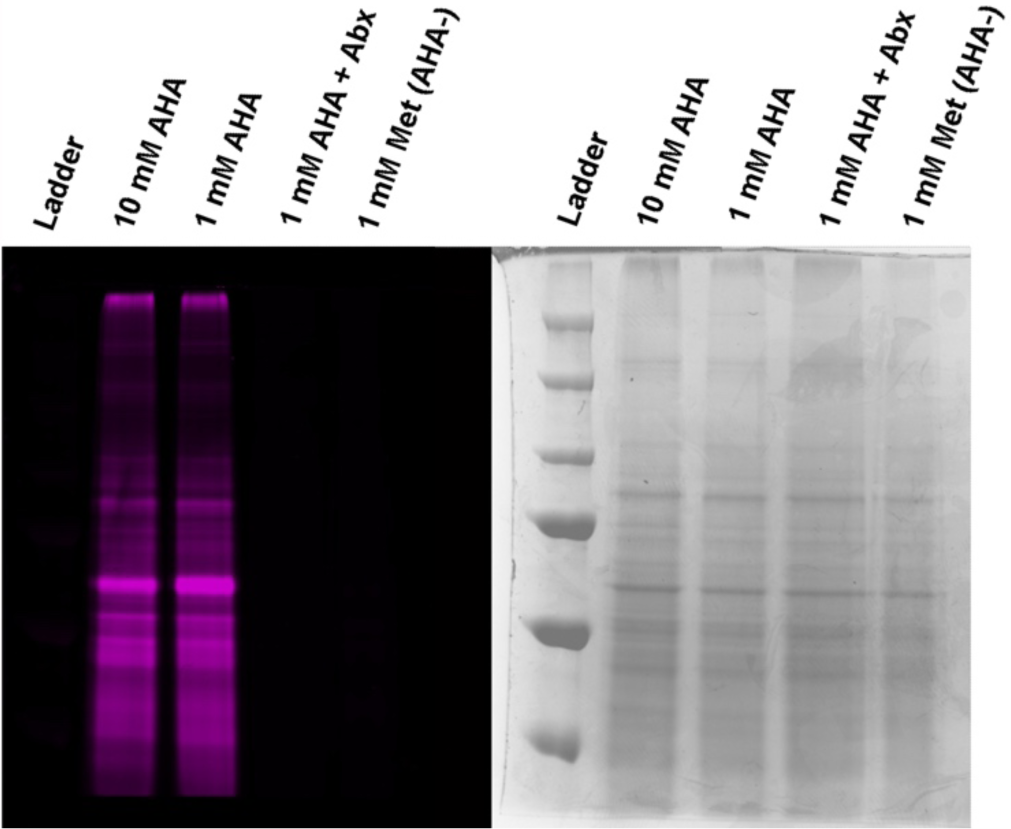
BONCAT labeling of *P. aeruginosa* is specific for AHA and translational activity. SDS-PAGE visualization of BONCAT labeling of laboratory cultures of *P. aeruginosa* PA14. Labeling is specific for AHA and inhibited by antibiotics (Abx = chloramphenicol, tetracycline, tobramycin).

**Supplementary Figure 3.**
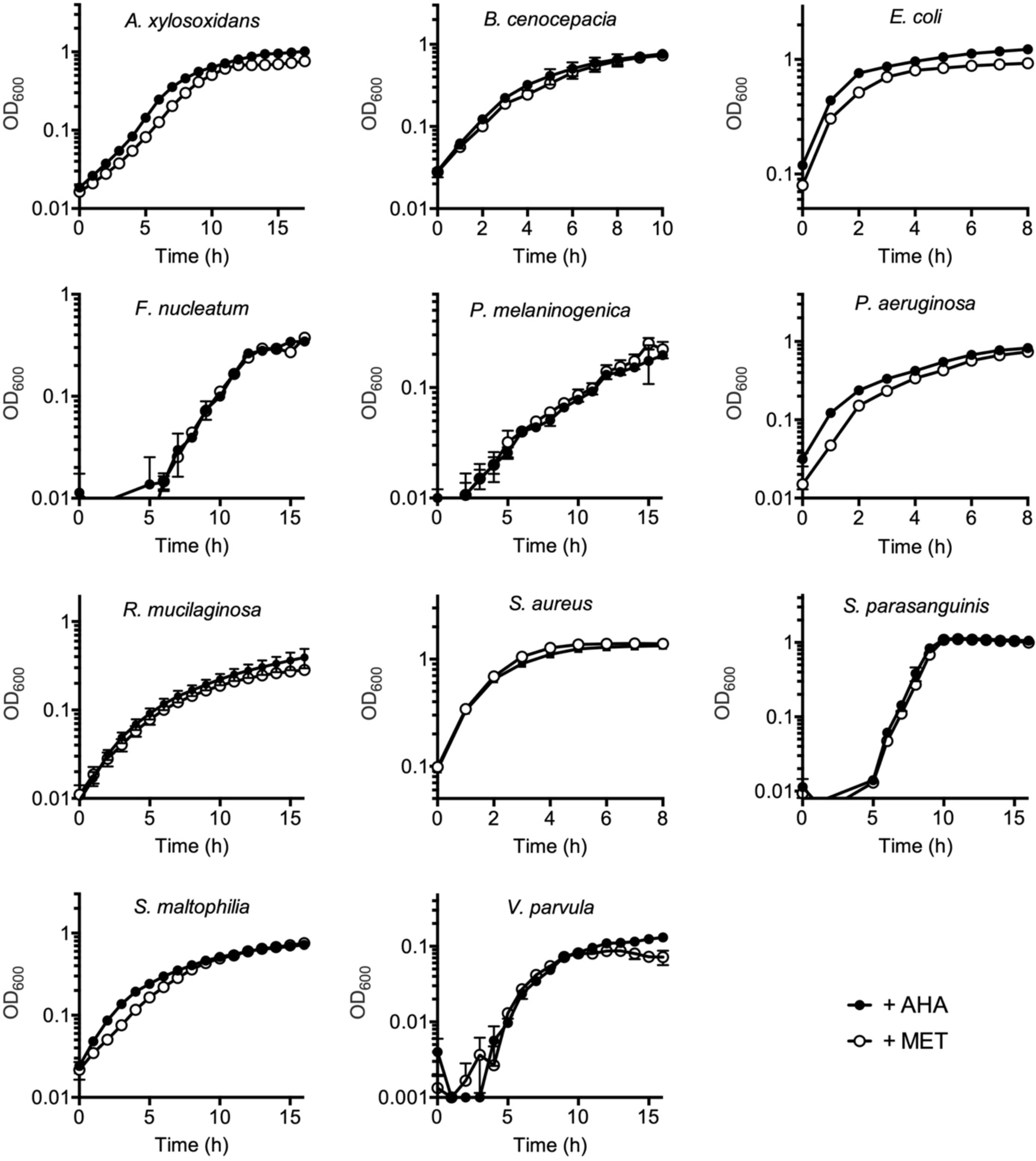
AHA-incubation has negligible effect on bacterial growth.

**Supplementary Figure 4.**
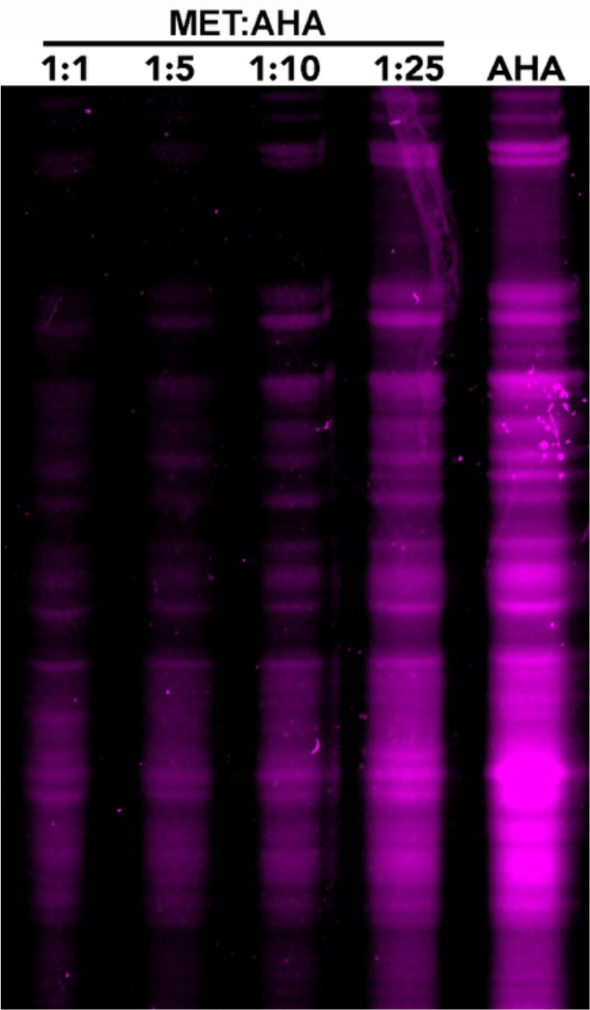
SDS-PAGE determination of MET:AHA ratio required for BONCAT labeling of *P. aeruginosa*. Cells were grown in varying concentrations of MET:AHA prior to labeling with Cy5-DBCO. Based on these profiles, a 1:10 ratio was selected for labeling of *in vitro* bacterial cultures and expectorated sputum samples.

**Supplementary Figure 5.**
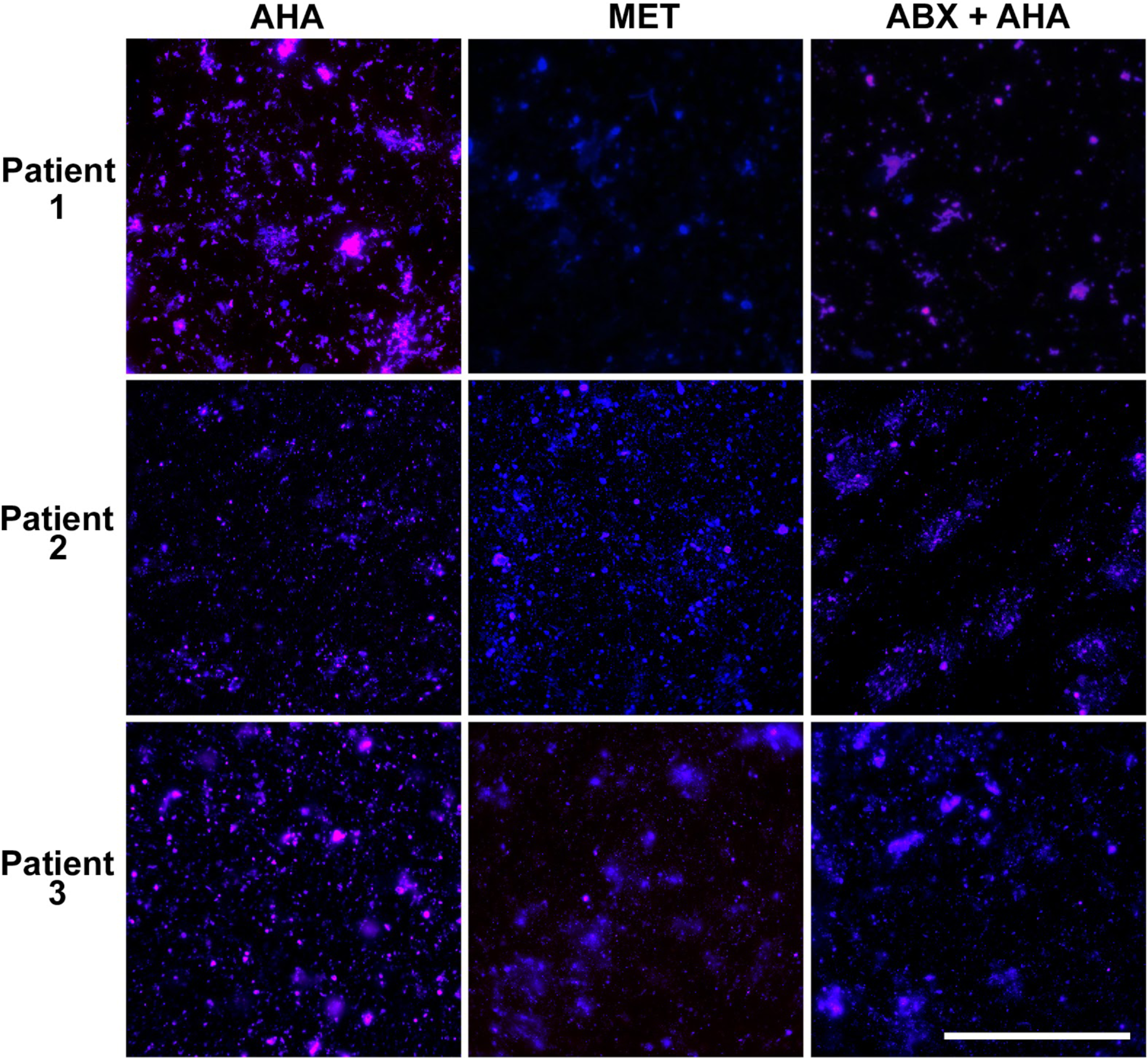
BONCAT labeling of sputum microbiota is specific for AHA. Expectorated sputum samples were supplemented with 6mM AHA and incubated for 3h prior to Cy5-DBCO labeling (Cy5; magenta) and counterstaining (SYTO64; blue). Negligible background fluorescence was observed in paired sputum samples incubated with methionine (MET). Incubation with antibiotics prior to AHA labeling (ABX+AHA) resulted in a moderate decrease in fluorescence that may reflect antimicrobial tolerance among airway microbiota. Bar = 100 µm. Abx = chloramphenicol, tetracycline, tobramycin.

**Supplementary Figure 6.**
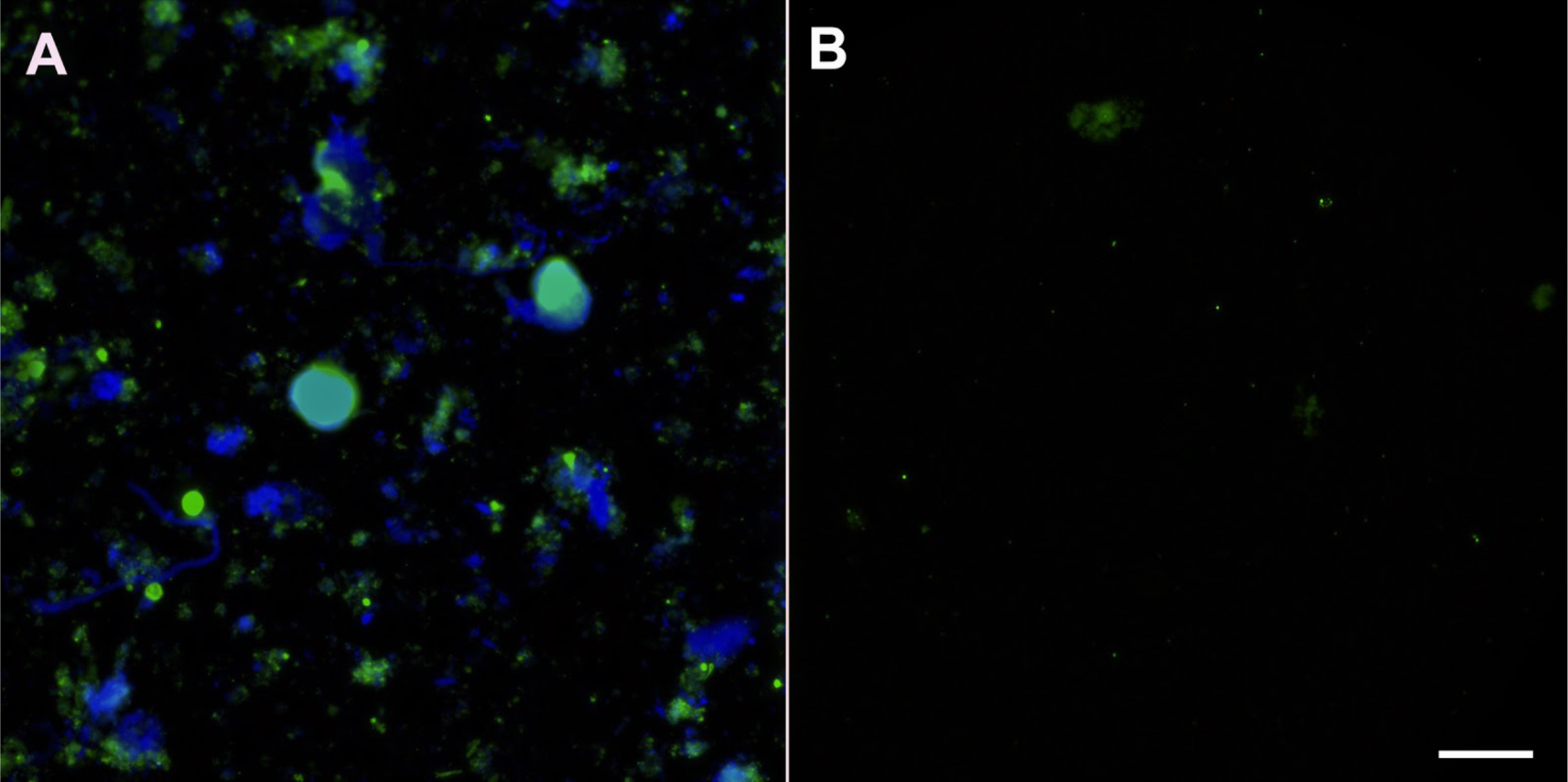
Validation of BONCAT-FACS. During sorting, the Cy5 label is subject to photobleaching; we therefore used immunostaining (anti-Cy5 antibody) and fluorescent microscopy to validate FACS sort integrity. Anti-Cy5 labeling of (a) “sort input” and (b) “sort positive” samples demonstrates that FACS effectively removes host cells and AHA-bacteria. As expected, all cells in the positive gate were anti-Cy5 reactive, confirming AHA uptake and translational activity among this bacterial subpopulation.

**Supplementary Figure 7.**
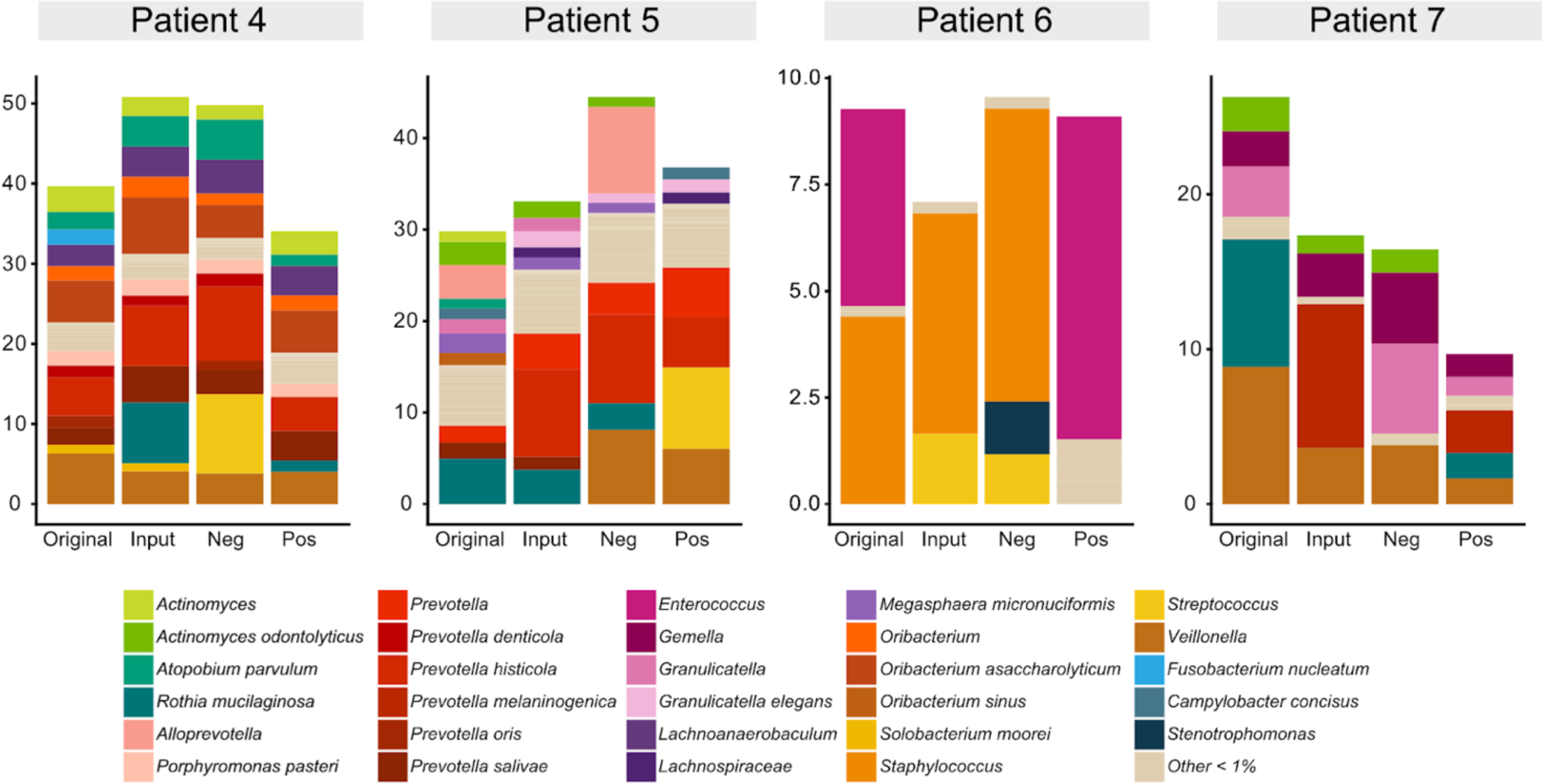
Low abundance community members are translationally active. Bacterial community membership of taxa with relative abundances less than 10%. Notable differences are observed between “sort positive” community composition (i.e. translational activity) relative to the “original” fraction.

**Supplementary Figure 8.**
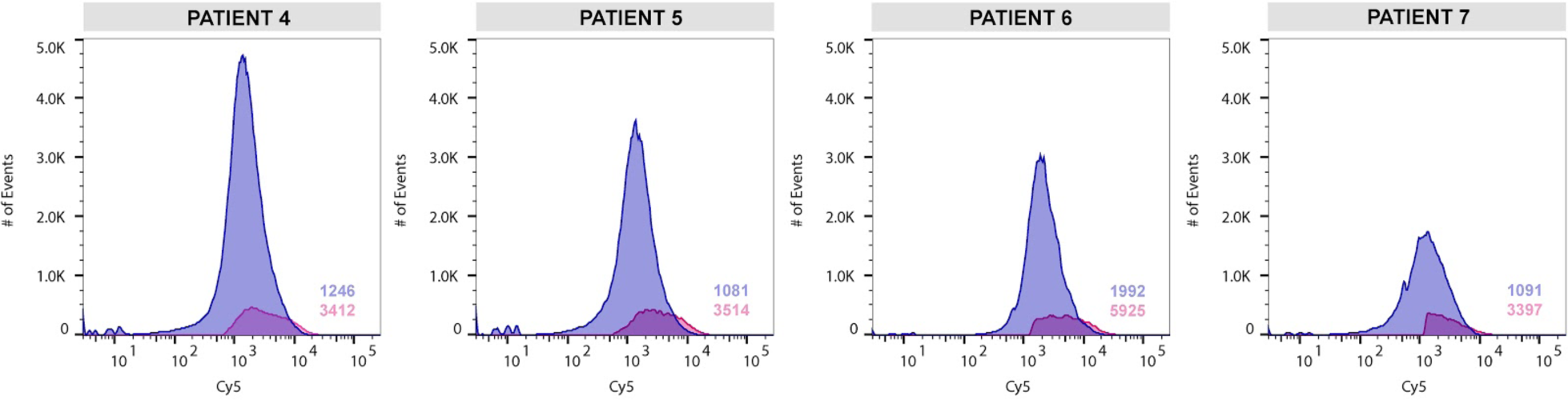
Confirmation of BONCAT Labeling. Flow cytometry histograms represent the number of events (cell counts) versus the log fluorescent intensity of Cy5 (blue = Cy5-population, magenta = Cy5+ population). Numbers in the lower right-hand corner represent the geometric means of Cy5 intensity. The Cy5+ population exhibited a higher geometric mean of fluorescent intensity validating AHA uptake and BONCAT labeling of the active subpopulation from CF sputum samples.

**Supplementary Figure 9.**
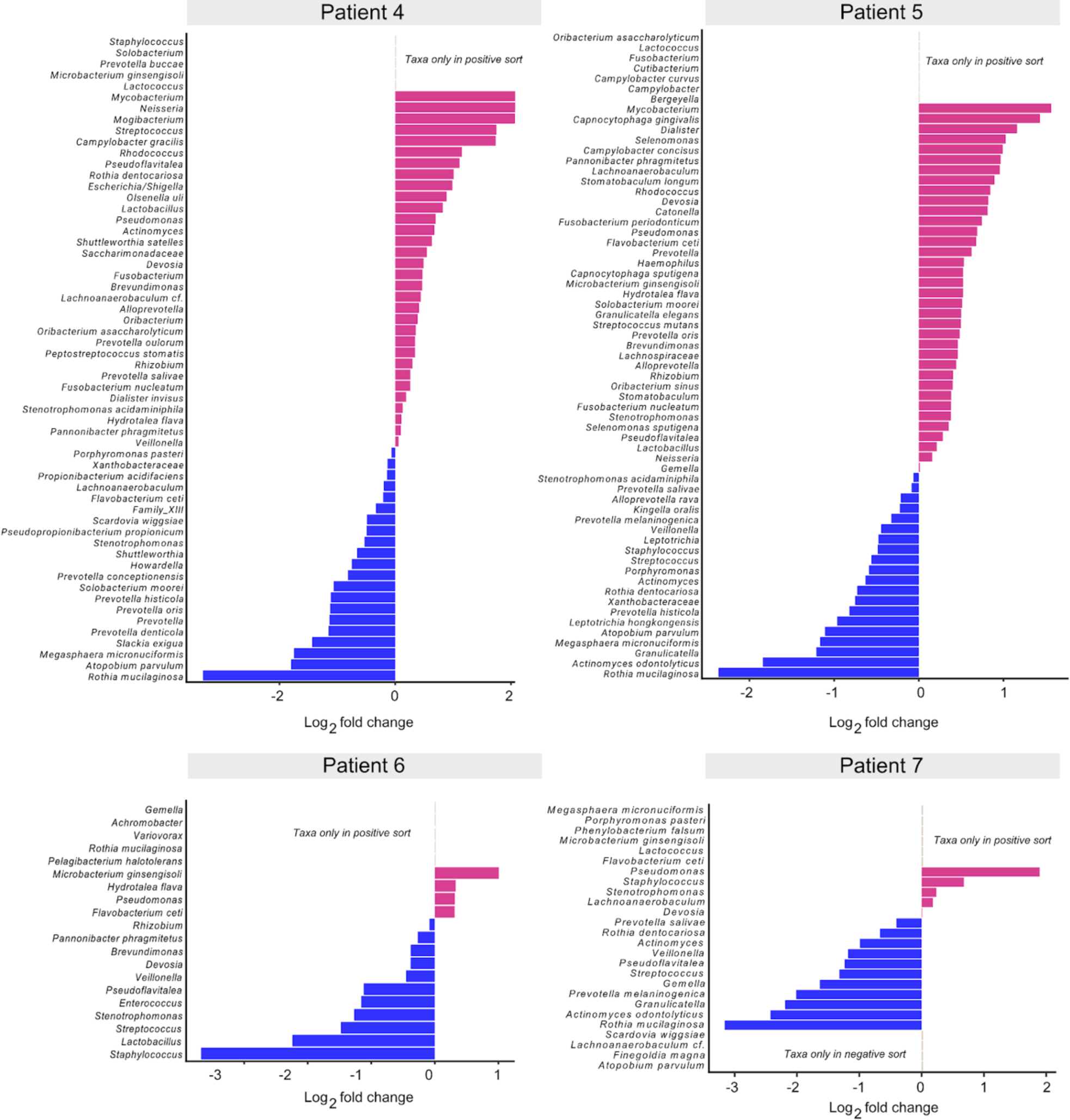
Log2 fold difference in taxon relative abundance between “sort positive” and “sort negative” fractions. Pink and blue bars indicate taxa that were increased in relative abundance in the positive and negative fractions, respectively.

**Supplementary Table 1.**
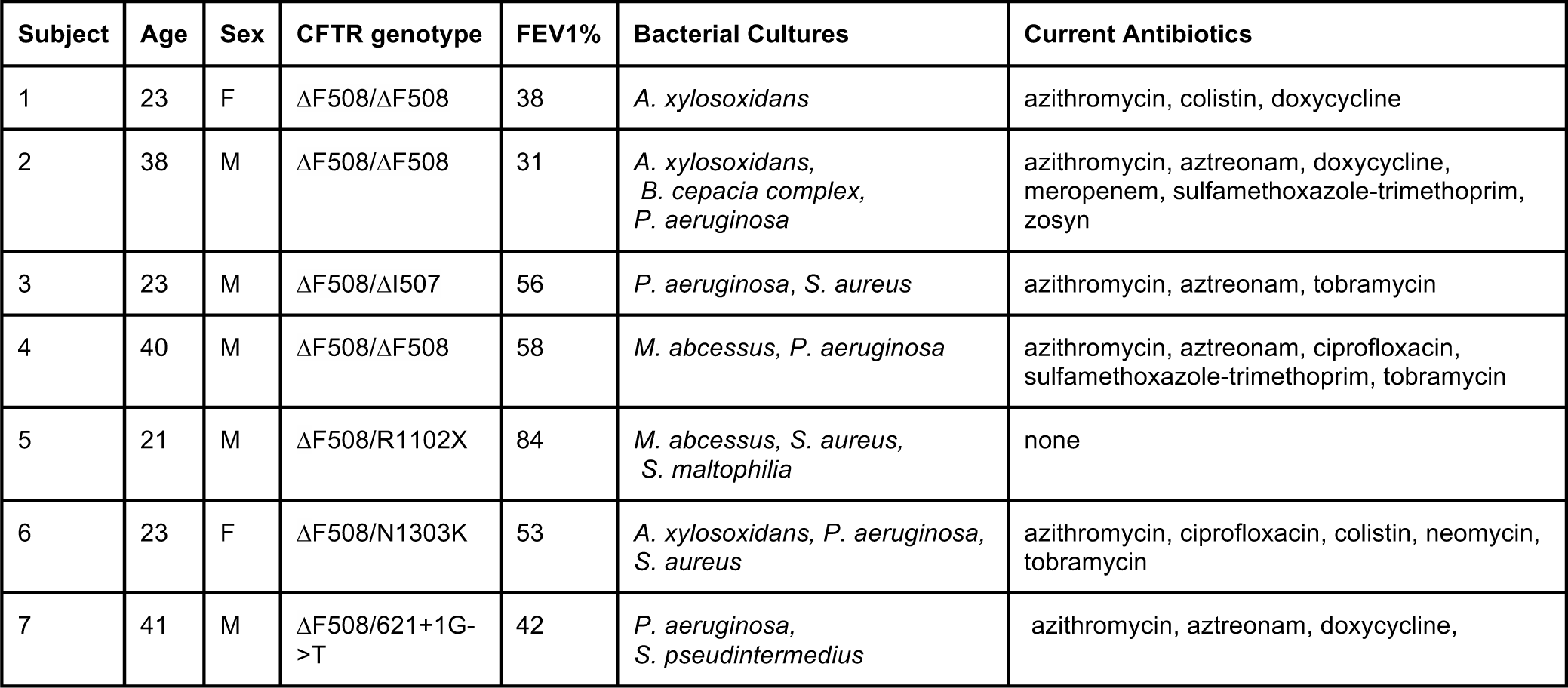
Patient clinical data.

**Supplementary Table 2.**
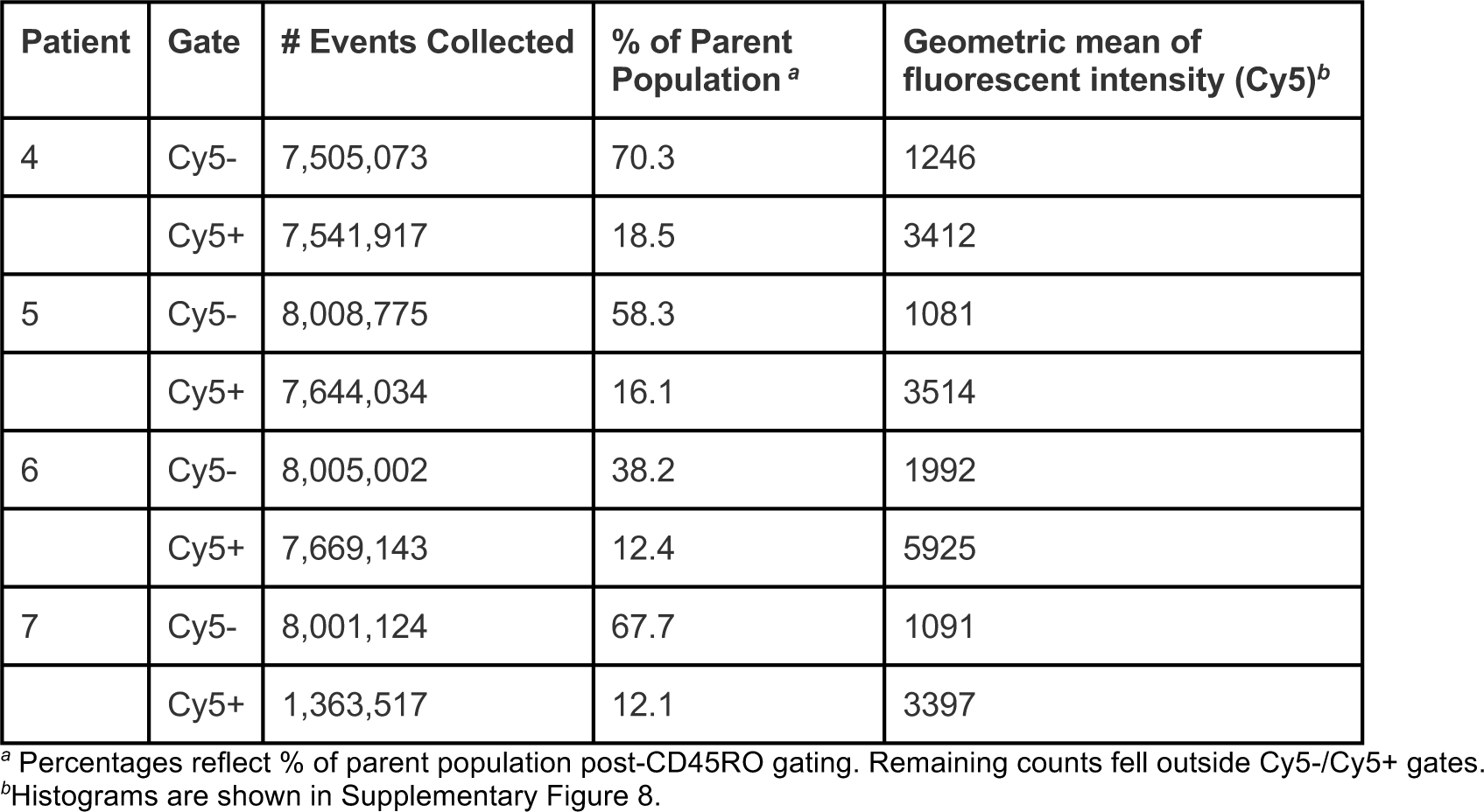
BONCAT-FACS summary.

